# Engineering an Enzyme for Direct Electrical Monitoring of Activity

**DOI:** 10.1101/678656

**Authors:** Bintian Zhang, Hanqing Deng, Sohini Mukherjee, Weisi Song, Xu Wang, Stuart Lindsay

## Abstract

Proteins have been shown to be electrically-conductive if tethered to an electrode by means of a specific binding agent, allowing single molecules to be wired into an electrical sensing circuit. This development opens the possibility of exploiting the remarkable chemical versatility of enzymes as sensors, detectors and sequencing devices. We have engineered contact points into a Φ29 polymerase by introducing biotinylatable peptide sequences. The modified enzyme was bound to electrodes functionalized with streptavidin. Φ29 connected by one biotinylated contact and a second non-specific contact showed rapid small fluctuations in current when activated. Signals were greatly enhanced with two specific contacts. Features in the distributions of DC conductance increased by a factor 2 or more over the open-to closed conformational transition of the polymerase. Polymerase activity is manifested by rapid (millisecond) large (25% of background) current fluctuations imposed on the DC conductance.

---

Proteins are widely assumed to be insulators. Reports of metallic conduction in bacterial wires^1^ and long range transport in protein multilayers^2^ were thought to be exceptions. We recently demonstrated that a number of proteins, chosen only for their *redox inactivity* (i.e. no electron transport function, no redox active centers) conduct very well if contacted by binding agents that can inject charge carriers into their interiors.^3^,^4^ This conductance is electron (or hole) mediated because measurements were made under potential control in conditions that eliminate Faradaic currents.^4^ The initial studies used proteins with multiple binding sites (antibodies, streptavidin) and found that conductance was sensitive to ligand binding,^4^ a finding that suggests that conductance might also be sensitive to conformational changes associated with enzyme activity. If so, this would pave the way to a whole new area of bioelectronics in which the remarkable chemical versatility of enzymes could be exploited by direct electrical measurements.^5^ Here, we demonstrate a technique for inserting binding sites into Φ29 polymerase, an enzyme that replicates a template DNA strand rapidly and accurately.^6^, ^7^ We also show that polymerase activity results in rapid conductance fluctuations.

## Engineering contacts

Design criteria were: (1) That contact points be remote from the active site of the enzyme; (2) That the sites not change relative spacing over the open to closed conformational transition; (3) That the spacing be as large as possible; (4) That the inserted sequences not change the isoelectric point of the enzyme. Measurements of polymerase activity served to test these objectives. In our first iteration, the Avitag peptide sequence was inserted near the N-terminus of the polymerase (Gen I). The lysine in this sequence is biotinylated using the BirA enzyme^8, 9^ allowing strong, and specific, binding to Streptavidin. Biotin-bound Streptavidin makes an excellent molecular wire^4^ and also serves to keep the Φ29 (with its seven surface cysteines) away from the metal electrodes. We made a second version (Gen II) with a second Avitag about 5 nm distant from the first, at a site in the (inactivated) exonuclease domain. A third version (Gen III) had both contacts as well as a flexible linker next to the N-terminal Avitag. Peptide sequences are given in Figure S1a, and visualizations of Generations I and III are shown in Figures 1a and b. Formation of the complex with streptavidin was confirmed with a protein gel, and the activity of the complex was demonstrated with a rolling-circle amplification assay (Figures S1b,c). The same assay was used to verify polymerase activity in the buffer used for STM measurements.

**Figure 1:**
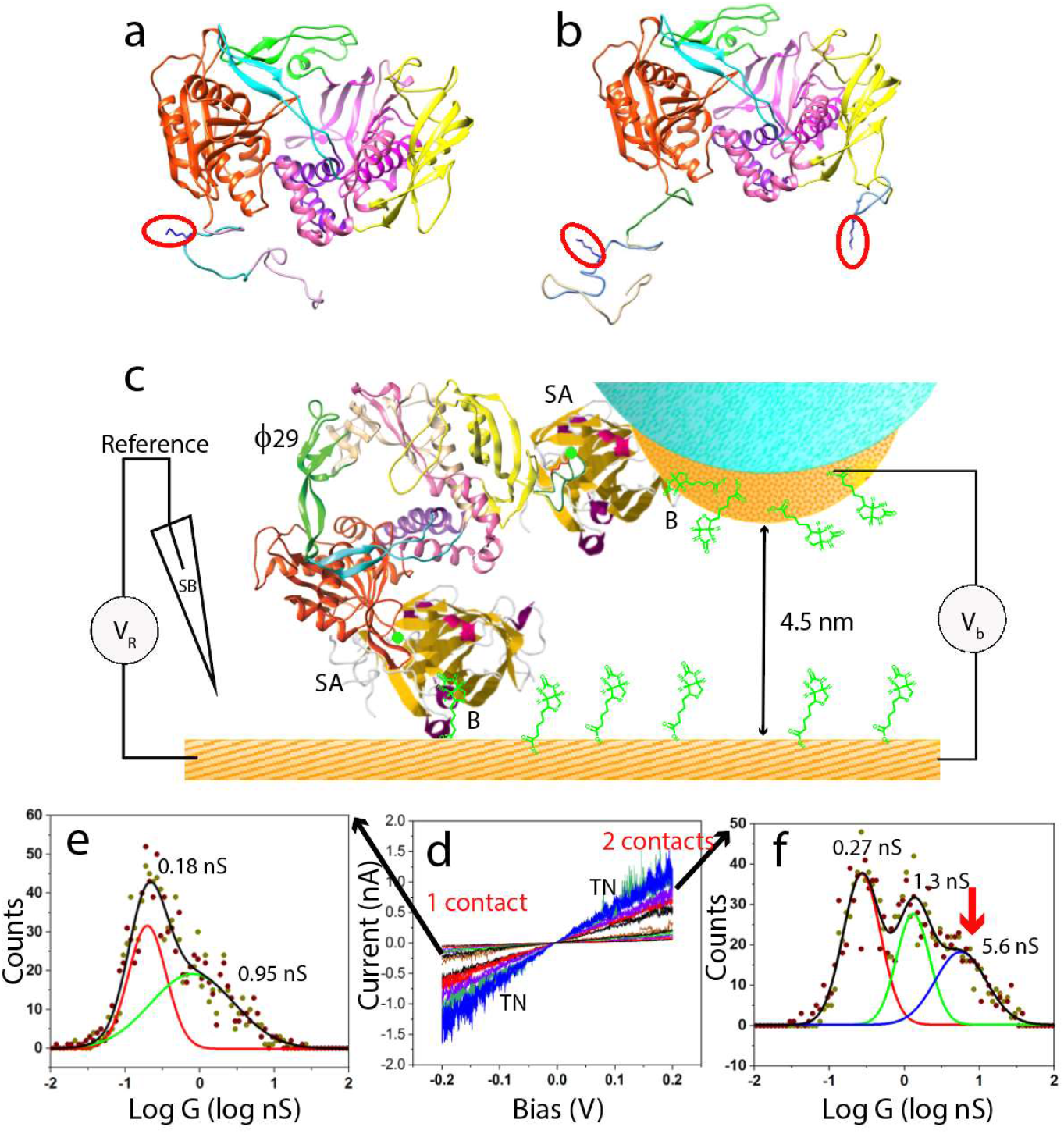
Conductance of polymerases with one and two biotinylated contact points. (a)Φ29 polymerase with a single Avitag at the N terminus (Gen I). Biotinylatable lysine is labeled by the red outline. (b) Φ29 polymerase with a second Avitag inserted between E279 and D280 and a flexible loop at N terminus (Gen III). (c) An STM probe is held ~4.5 nm above a conducting substrate, immersed in electrolyte and under potential control via a salt bridge (“SB”) to an Ag/AgCl reference. Electrodes functionalized with thiolated biotin (“B”) capture streptavidin molecules (“SA”) which trap a biotinylated polymerase (“Φ29”). (d) Typical current-voltage curves (trace and retrace are superimposed). Conductances for individual molecules are obtained from the slopes of these traces. “TN” indicates region of contact-field induced fluctuations. A doubly biotinylated polymerase has a new high conductance feature at ~6 nS in the conductance distribution (red arrow in f) not present in the singly-biotinylated molecule (e). The largest uncertainty in the fitted peaks is ± 0.05 in log G, corresponding to about ±0.12G in G.

## Scanning Tunneling Microscope Conductance Measurements

Measurements were made using an electrochemical scanning tunneling microscope (Pico STM, Agilent) with insulated palladium (Pd) probes^10^ and a Pd substrate, both held under potential control using a salt-bridged reference electrode (Figure. 1c). Electrodes were modified with streptavidin using either a thiolated biotin (SH-biotin) or thiolated streptavidin, and then incubated with a solution of the biotinylated polymerase (Methods). Measurements were made in a reaction buffer containing MgCl2 and tris(2-carboxyethyl)phosphine) (TCEP) to prevent polymerase oxidation. Nucleotide triphosphates were added to activate the polymerases. Current-voltage (IV) characteristics were measured using a fixed Z gap (no servo control) which remained constant to within about 0.1nm over ~ 1 minute, as determined by tunnel-current measurements. Drift in the X-Y plane cannot be measured so accurately, but the contact point with a target molecule clearly changes over time. The bias was swept between −0.2 and +0.2V and back again at a rate of 1V/s. After 1 minute, the gap was returned to the set-point value and the cycle was repeated to obtain further IV sweeps. 80% of these sweeps were linear and reproduced exactly on reversing the sweep direction (Figure 1d). The gradients of these sweeps were used to compile conductance distributions that reflect different types of contacts (Figures 1e,f).

The molecular junctions (Figure 1c) were assembled by coating the electrodes with streptavidin, using thiolated streptavidin or wild-type streptavidin in combination with biotinylated electrodes (Methods). Conductance through the streptavidin alone is only observed when the gap is less than 3.5 nm. When biotinylated Φ29 is added to the liquid cell, high conductance is observed out to a 4.5 nm gap (Figure S2). Unexpectedly, the monobiotinylated Gen I polymerase gave 2 conductance peaks (Figure 1e). The first peak (at ~0.2 nS) is characteristic of one specific and one non-specific contact.^4^ The additional peak may reflect interactions between surface cysteines on the polymerase and the electrodes (Figure. 3b). The bis-biotinylated Gen III displays three conductance peaks (Figure 1f), the highest of which (~5.6 nS) is characteristic of a bridge formed by binding of specific ligands.^4^ Note that connection via biotinylated electrodes gives higher conductance than direct attachment via thiolation of surface residues (Figure. S3a). We conclude that the highest conduction peak corresponds to conductance through the polymerase molecules. Accordingly, we next explored the conformational dependence of the electronic conductance.

The active domain of the polymerase resembles a human hand, with a thumb subdomain that holds the DNA, a palm subdomain that contains the catalytic site, and a moving fingers subdomain that closes around the complex of DNA template once the correct complementary nucleotide triphosphate (dNTP) is bound. The enzyme is normally ‘open’ and remains so after binding DNA containing a primer strand and template strand with a 5’ −overhang. Once the correct dNTP is bound, the fingers close to complete the reaction, opening again only for long enough to bind the next complementary dNTP.^11^ This transient opening can be suppressed by using non-hydrolyzable dNTPs (NH-dNTPs) in which a carbon replaces an oxygen in the triphosphate.^12^

We repeated measurements of the conductance distributions: (a) With a saturating (1μM) concentration^13^ of a single-stranded template with a 15 basepair hairpin primer (Figure S4). (b) With the template-bound polymerase in the presence of a saturating concentration^13^ (1mM) of dNTPs. (c) In the presence of a saturating concentration^13^ (1mM) of NH-dNTPs. The corresponding conductance distributions are shown in Figures 2a, b and c. The distribution in the presence of bound template (Figure 2a) is almost identical to the distribution in the absence of template (Figure 1f - uncertainties in these fits are discussed in the caption). On addition of dNTPs there are large shifts in the conductance peaks (Figure 2b - 1.3×, 2.3× and 2× for peaks 1,2 and 3). Locking the polymerase in the closed form changes the peak positions a little more (Figure 1c, 1.8×, 3.2× and 2.3×) with a notable sharpening of the third peak (see Figure S5 for replication of this sharp peak). Thus we conclude that the open to closed transition of the polymerase is accompanied by large changes in conductance.

**Figure 2:**
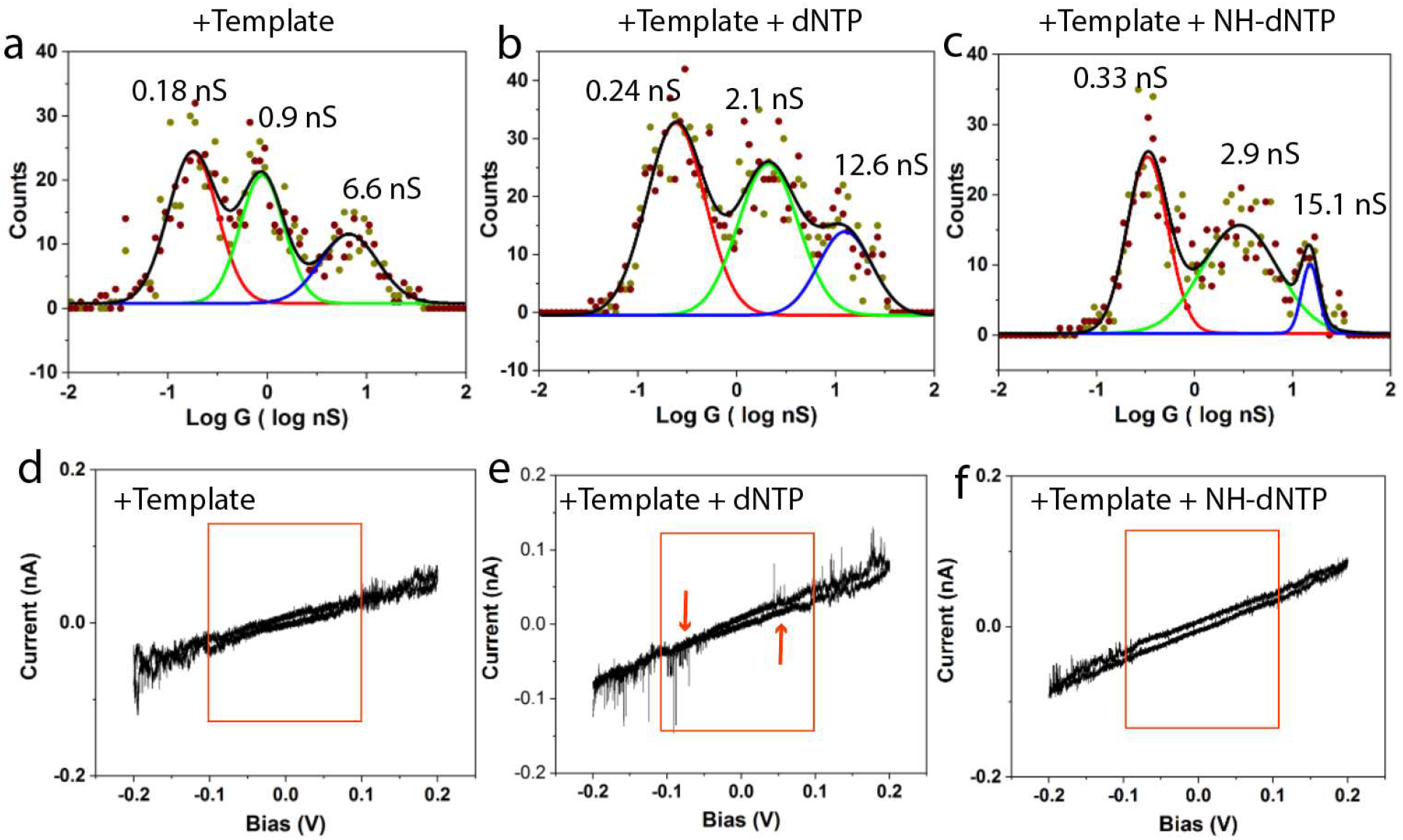
The open to closed transition changes polymerase conductance. (a) Distribution in the absence of dNTPs but with bound template. The polymerase is largely open. (b) With dNTPs added (mostly closed) the distribution changes dramatically. (c) Conductance distribution for a polymerase locked in the closed conformation with non-hydrolyzable dNTPs. (d,e,f) In the inactive state (−dNTP d, or + NH-dNTP f) the IV curves are noise free in the bias range below ±100mV (red box). The active polymerase (e) shows noise spikes on the IV curve in this otherwise quiet region (red arrows).

**Figure 3:**
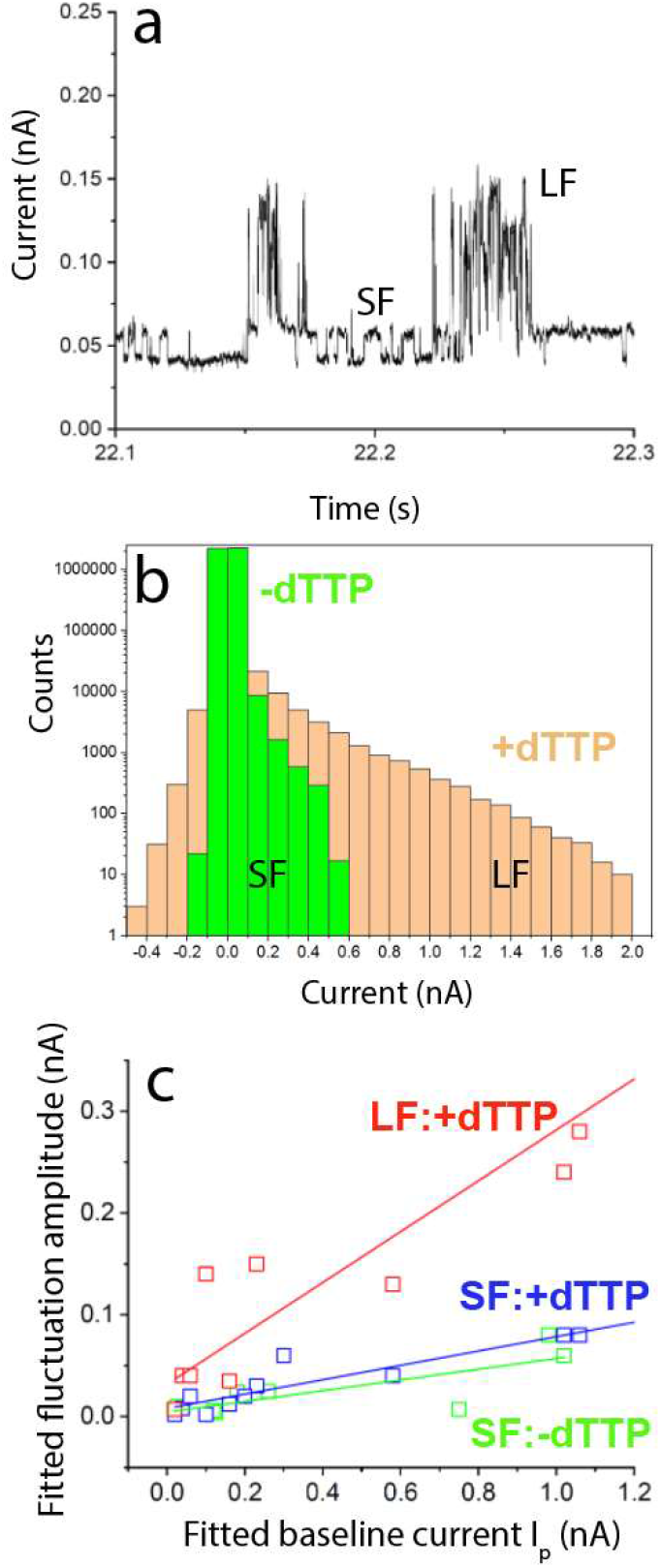
Characterizing the signals generated when polymerase is activated. (a) An expanded portion of the noise signal measured from an activated polymerase (A_10_ template) showing two distinct noise components - large fluctuations (LF) and small fluctuations (SF). (b) Distributions of noise signal amplitudes from the active (+dTTP) and inactive (-dTTP) polymerase in runs of about the same background current. (c) The peak noise amplitudes, I_I_ (LF) and I_S_ (SF), depend on the baseline current I_P_. I_l_ (red) and I_S_ (blue) are plotted vs the associated value of baseline current (I_P_). Only SF are observed in inactive (−dTTP) polymerases (Green). Similar results for other template sequences are shown in Figure S10.

These measurements were taken in the presence of Mg^2+^ so the polymerase is catalytically active in the presence of both template DNA and dNTPs. This is marked by additional noise, as shown in samples of the IV curves for the three cases in Figures 2d,e and f. The inactive polymerase (Figures. 2d,f) is essentially noise-free in the bias region below ± 100mV, as reported for other proteins.^4^ (Above 100mV, the electric field at the contact points induces telegraph noise - the current at a fixed bias switches between two distinct levels.^4, 14^) However, when the polymerase is active (Figure 2e) large spikes are also observed in the bias region below ± 100 mV.

## Noise Measurements

These observations indicate that rapid polymerase activity can be followed by recording current versus time (I(t)) at a bias below 100mV in the presence of template-bound Φ29, Mg^2+^ and dNTPs. We set a gap of 2.5 nm under servo control, opened the servo, increased the gap to 6 nm, then brought the tip down to 4.5 nm and recorded current for 60 s at a bias of 50mV. Typically, no current was recorded for the first 10-20 s, after which a contact formed and an I(t) curve was obtained. Contacts were formed with molecules in >50% of these “fishing” attempts. The currents jumped suddenly on contact with the molecule, but then changed substantially as the contact point drifted. A typical current-time trace is shown in Figure S6a. This variation in current vs. time was shown to be a consequence of the drift in the contact point - the distribution of currents taken in an I(t) curve replicates the distribution measured by taking IV curves from many different contact points.^4^

Telegraph noise is clearly visible in I(t) traces taken with activated polymerases. Figure S7 shows examples for the Gen I, single contact polymerase. However, these data also illustrate how difficult it is to quantify the noise in the presence of a rapidly varying background current.

We removed the background using an asymmetric least squares (ALS) fit.^15^ The ALS accurately follows the background without distorting the noise signals (Figure S8). The I(t) trace shown in Figure S6a was obtained with a dA10 template (Figure S4b) in the presence of dTTP. The raw data are shown as black points, largely obscured by the ALS fit (red). The subtracted signal, corresponding to the fluctuations, is shown in Figure S6b. When the same procedure is applied to a trace taken at about the same current in the absence of dTTP (Figures S6c and d) it is clear that noise is also present in the −dTTP control. However the noise is much smaller in amplitude. A closer examination of the signals reveals two distinct levels of telegraph noise, as shown in Figure 3a and labeled “SF” (small fluctuations) and “LF” (large fluctuations). Noise-amplitude distributions for the traces in Figures S6b and d are shown in Figure 3b. The SF appear in all measurements whereas the LF are only present when the polymerase is active. Quantifying this qualitative observation is complicated because the absolute amplitude of the fluctuations depends on the background current, and this current varies over a run. However, the run-to-run variations are generally much larger, so we obtained an approximate measure of the relative fluctuation size as follows: For each molecule measured, we binned the ALS fitted baseline currents as shown by the examples in Figures S9a,c. Many of these distributions could be fitted by a Gaussian (red curve). Many could not - for example the background can jump between two or three levels. In these cases we fitted the largest peak that was clear of the background. The peak of the fitted Gaussian, I_p_, was then used to characterize the baseline for that run. Example of the binned noise signals are given in in Figures S9b,d. To characterize these we used a double exponential distribution:

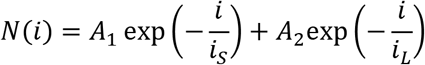

where *i* is the current in a given bin of the current distribution. In the case of the experiments in which dTTP was withheld (e.g., Figure S9b), the fits all converged to a single exponential (*i_S_* = *i_L_*). For recordings with dTTP present, most of the fits converged on the double exponential distribution with *i_S_*<*i_L_* (4 of 13 molecules showed the small peaks only in the presence of dTTP). The results are summarized in Figure 3c. Most activated molecules (+dTTP) showed both large (red) and small (blue) fluctuations. The controls (-dTTP) showed only small fluctuations (green), essentially equal to the small fluctuations also seen in the active polymerases. There is an approximately linear relation between background current and fluctuation amplitude as shown by the three linear fits. For the large fluctuations, characteristic of active polymerases, a typical (1/e value) of current is *i_L_* = (0.25 ± 0.026)*i_p_* For the small fluctuations, present in both active an inactive polymerases, a typical value is *i_S_* = (0.06 ± 0.01)*i_p_*. Thus the active state can be identified by the presence of fluctuations that are about 25% of the baseline current, while fluctuations in the inactive state are about 6% of the baseline current. Not all polymerase molecules contacted were active as indicated by the lack of large fluctuations in 4 of the 13 molecules studied. On the other hand, none of the eight −dTTP control runs showed large fluctuations (Table 1).

**Table 1:** Occurrence of large fluctuations (fraction of measured molecules) for various experimental conditions. NH = non-hydrolyzable dNTPs. Large fluctuations are identified by a fit converging on a two component distribution of amplitudes.

| Experimental Conditions | Control/Active | Fraction Large Fluctuations |
| --- | --- | --- |
| (dA)<sub>10</sub> +dNTPs +Mg<sup>2+</sup> | A | 0.69 |
| (dA)<sub>10</sub> +dATP, + dCTP, +dGTP, -dTTP, + Mg<sup>2+</sup> | C | 0 |
| (dC)<sub>10</sub> +dNTPs +Mg<sup>2+</sup> | A | 0.52 |
| (dA)<sub>10</sub>(dC)<sub>10</sub> +dNTPs +Mg<sup>2+</sup> | A | 0.44 |
| (dATC)<sub>5</sub> +dNTPs +Mg<sup>2+</sup> | A | 0.52 |
| (dA)<sub>10</sub>(dC)<sub>10</sub> - dNTPs + Mg<sup>2+</sup> | C | 0 |
| (dA)<sub>10</sub>(dC)<sub>10</sub> +dNTPs - Mg<sup>2+</sup> | C | 0 |
| (dA)<sub>10</sub>(dC)<sub>10</sub> +dTTP -dGTP, + Mg<sup>2+</sup> | C | 0.32 |
| (dA)<sub>10</sub>(dC)<sub>10</sub> NH-dNTPs +Mg<sup>2+</sup> | C | 0 |
| No template +dNTPs +Mg<sup>2+</sup> | C | 0 |

This analysis was repeated using data obtained in 38 runs with a d(ATC)_5_ template (Figure S4d) and 25 runs with a d(C)_10_ template (Figure S4a). The results are summarized in Figures S10a and b. The fitted amplitude distributions for the large fluctuations (LF) show considerable variation, but the trends observed for d(A)_10_ (Figure 3c) are reproduced well with *i_L_* = 0.27(±0.03)I_p_ and *i_s_* = 0.04(±0.01)I_p_ for d(ATC)_5_ and *i_L_* = 0.32(±0.03)I_p_ and *i_s_* = 0.05(±0.007)I_p_ for d(C)_10_.

In order to confirm the association of large fluctuations (amplitudes > 25% of the baseline current) with polymerase activity we carried out experiments with several DNA templates (Figure S4) in different conditions. Active (“A” in Table 1) polymerases were measured in the reaction buffer (1mM phosphate buffer, pH=7.4, 4 mM TCEP, 10mM MgCl2 with 1 mM dNTPs and 1μM template). In the control experiments (“C” in Table 1), one essential ingredient was withheld. In addition, we carried out measurements using nonhydrolyzable dNTPs.

Fluctuations were analyzed as described above.

Clearly, withholding any one of the ingredients critical to polymerase function abolishes the large fluctuations. An interesting exception was (dA)_10_(dC)_10_ in the presence of dTTP only. We expected that the polymerase would reach the end of the A tract and then stall for want of the missing dGTP nucleotide, but about a third of the molecules appear to be active for at least some part of the recording. We noted that the telegraph noise obtained from the (ATC)_5_ and C_10_ templates was generally more regular in time and amplitude than was the case for the A10 template (Figures 4a,b,c) and a denaturing gel of the products of polymerization (Figure 4d) clearly shows incomplete transcription of the A_10_ template (red arrow, lane 7). Thus it seems likely that the polymerase falls off the A homopolymer tract, allowing another template to bind, and accounting for the activity seen for the A_10_C_10_ template in the absence of dGTP.

**Figure 4:**
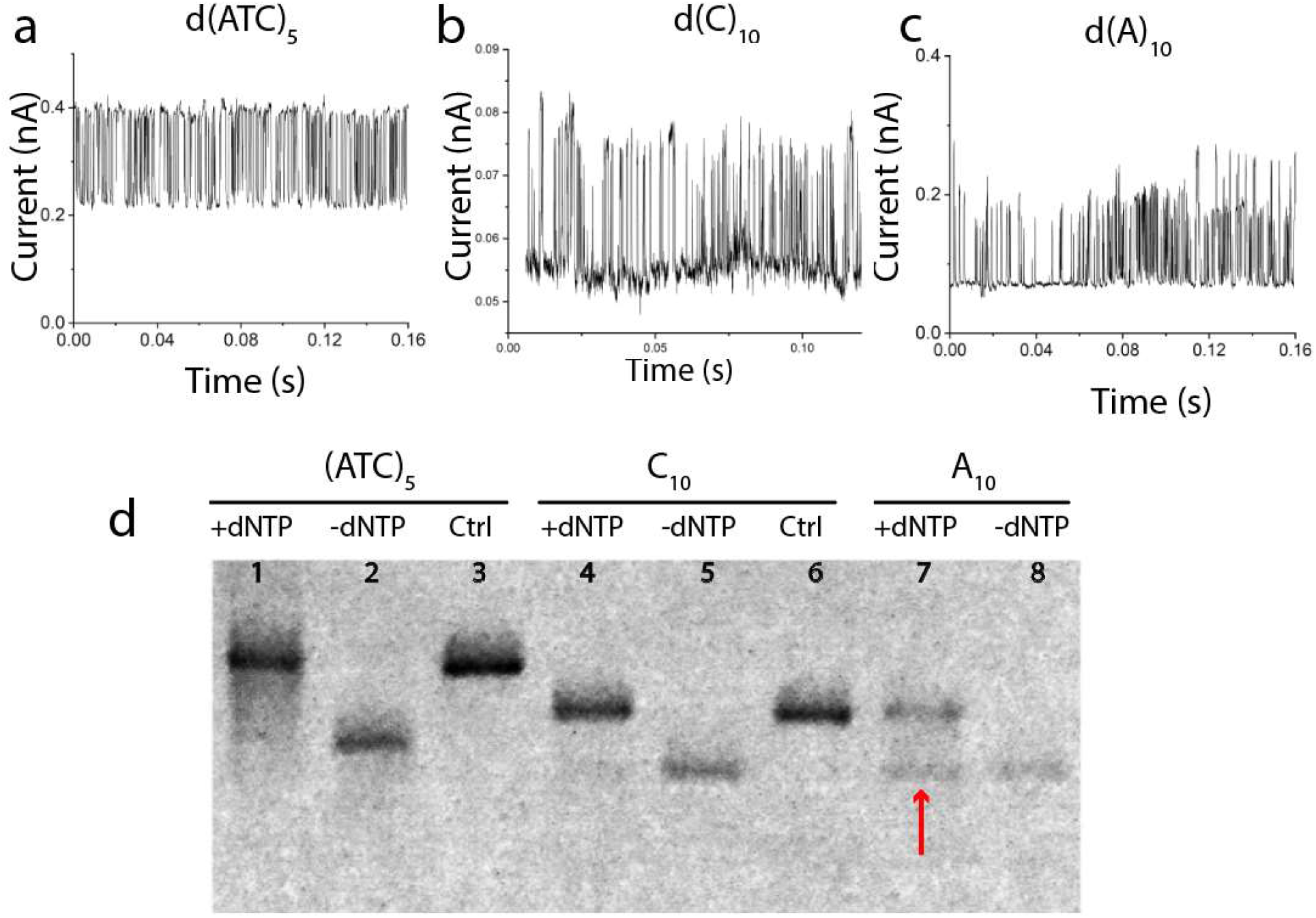
Telegraph noise signals reflect the ease with which a template is processed. Typical telegraph noise signals (selected from regions of constant baseline current) for d(ATC)_5_ and dC_10_ are fairly uniform in amplitude and in the period between signal features. Signals for dA10 are typically more irregular (c). A denaturing gel (d) shows that the d(ATC)_5_ and dC_10_ templates are completely converted to full length molecules (the Ctrl lanes are for synthesized fully extended molecules). This is not the case for the dA_10_ (red arrow) for which polymerization is only partially successful (The C_10_ control serves as a length control for this molecule also).

## CONCLUSIONS

Engineering two contact points into a polymerase yields features in the conductance distribution that are approximately 3 to 10 times larger than those observed with only one engineered contact and a second, non-specific, contact. The conductance of the complex of streptavidin and doubly biotinylated Φ29 is further increased if biotin is used to anchor the streptavidin to the electrodes instead of thiolated surface lysines. There are large changes in the conductivity distribution as the polymerase undergoes the open to closed transition. In addition, polymerase activity is marked by rapid noise spikes that have an amplitude of about 25% (or more) of the background current, distinct from the smaller (6% of background) signals present in both active and inactive polymerase. The difficulties caused by the unstable STM contact are mitigated when a solid state junction is used^14^ and we hope to gain further insight into the incorporation kinetics of nucleotides using these devices.

## METHODS

### Recombinant Φ29 DNA polymerase constructs with inserted Avitag

The starting enzyme was a Φ29 DNA polymerase, rendered exonuclease-deficient with D12A and D66A mutations. A Q5 site-directed mutagenesis kit (NEB) was used to insert the Avitag DNA sequence into a pET15b plasmid containing the mutant polymerase gene. The equivalent inserted peptide sequence is shown in blue, with flanking linker sequences in yellow, in Figure S1a. The epsilon-amine of the central lysine (K, marked by red arrow on Fig. S1a) was biotinylated using the BirA enzyme.^8^ Three generations of modified enzyme were tested. The first (Gen 1) was biotinylated only at the N terminus of the protein. The second (Gen II) contained a second Avitag ~5nm from the N terminus between E279 and D280. This second site is located in the deactivated exonuclease domain and was chosen because its position does not change with respect to the N terminus over the open to closed transition.^16, 17^ The third (Gen III) contained an additional flexible linking sequence (GNSTNGTSNGSS) adjacent to the N-terminal Avitag to allow for greater flexibility in contact geometry. Biotinylation was verified with a SDS-PAGE gel analysis of the free-and streptavidin bound polymerases. Figure S1b shows the increase in molecular weight that occurs as streptavidin binds to the Gen III polymerase (yet higher molecular weight features are likely polymer aggregates of alternating polymerase and streptavidin). The activity of the modified polymerases was verified in vitro using a rolling circle amplification assay (Figure S1c). Further details are given in the caption for Figure S1a.

### Functionalizing substrates and STM probes

Palladium substrates were prepared by evaporating a 200 nm palladium film onto a silicon wafer using an electron-beam evaporator (Lesker PVD 75), with a 10 nm titanium adhesion layer. The substrates were treated with a hydrogen flame immediately before functionalizing and then immersed in solutions of thiolated streptavidin (ProteinMods) or thiolated biotin overnight. The thiolated biotin was prepared as described elsewhere^4^ and dissolved in freshly degassed pure ethanol to a final concentration of 50 μM. 1 μM thiolated streptavidin solutions in 1 mM PB buffer were used for substrate functionalization. All the buffers and solutions were prepared in Milli-Q water with a conductivity of 18.2 MΩ. For all measurements, the 1 mM PB buffer (pH 7.4) was degassed with argon to avoid interference from oxygen. The polymerization buffer was 1mM phosphate buffer, pH=7.4, 4 mM TCEP, 10mM MgCl_2_ with 1 mM dNTPs and 1 μM template (activity in this buffer was verified with a rolling circle amplification assay). Substrate functionalization was characterized by Fourier transform infrared (FTIR) spectroscopy (Figure. S11). STM probes were etched from a 0.25mm Pd wire (California Fine Wires) by an AC electrochemical method. To avoid current leakage, probes were insulated with high-density polyethylene following the method described earlier for gold probes.^10^ Each probe was tested by STM in 1 mM phosphate buffer (pH 7.4) at +0.5 V bias to ensure the leakage current was <1pA. For functionalization, the probe was immersed in ligand solutions for 4 h or overnight, then rinsed with water, dried with nitrogen gas, and used immediately. Cyclic voltammetry was used to check that the potential regions used (+50 to −50mV vs Ag/AgCl, 10 mM KCl) were free from Faradaic currents in the presence of the various components of the assembly (Figures S12-14).

### STM Measurements

STM measurements were carried out on a PicoSPM scanning probe microscope (Agilent Technologies), using a DAQ card (PCI-6821 or PCIE-7842R, National Instruments) for data acquisition. The Teflon liquid cell was cleaned with Piranha solution and sonicated in Milli-Q water (Note that Piranha solution is highly corrosive and must be handled with extreme care). An Ag/AgCl reference electrode with a 10 mM KCl salt bridge was connected onto the substrate. The probe was initially engaged at a 4 pA setpoint current with a bias of −0.2 V and left to stabilize for 2 h before measurement. IV sweep and I(t) measurements are described in detail in reference ^4^.

## ASSOCIATED CONTENT

Online supporting Information: Figures S1 to S14.

## Supporting information

SI for Engineering an Enzyme

## Notes

The authors declare the following competing financial interest(s): SL is a cofounder of a company with interests in this the subject matter of the present paper. SL, BZ and HD are named as co-inventors on patent applications.

## ACKNOWLEDGMENT

This work was supported by grants HG006323 and HG010522 from the National Human Genome Research Institute, by Recognition AnalytiX Corp and the Edward and Nadine Carson Endowment.

## Notes

#### Summary of Updates

Paper has been shortened and better gel data obtained on the activity of the modified enzyme.

