## Supplementary material for "Engineering an Enzyme for Direct Electrical Monitoring of Activity": SI for Engineering an Enzyme


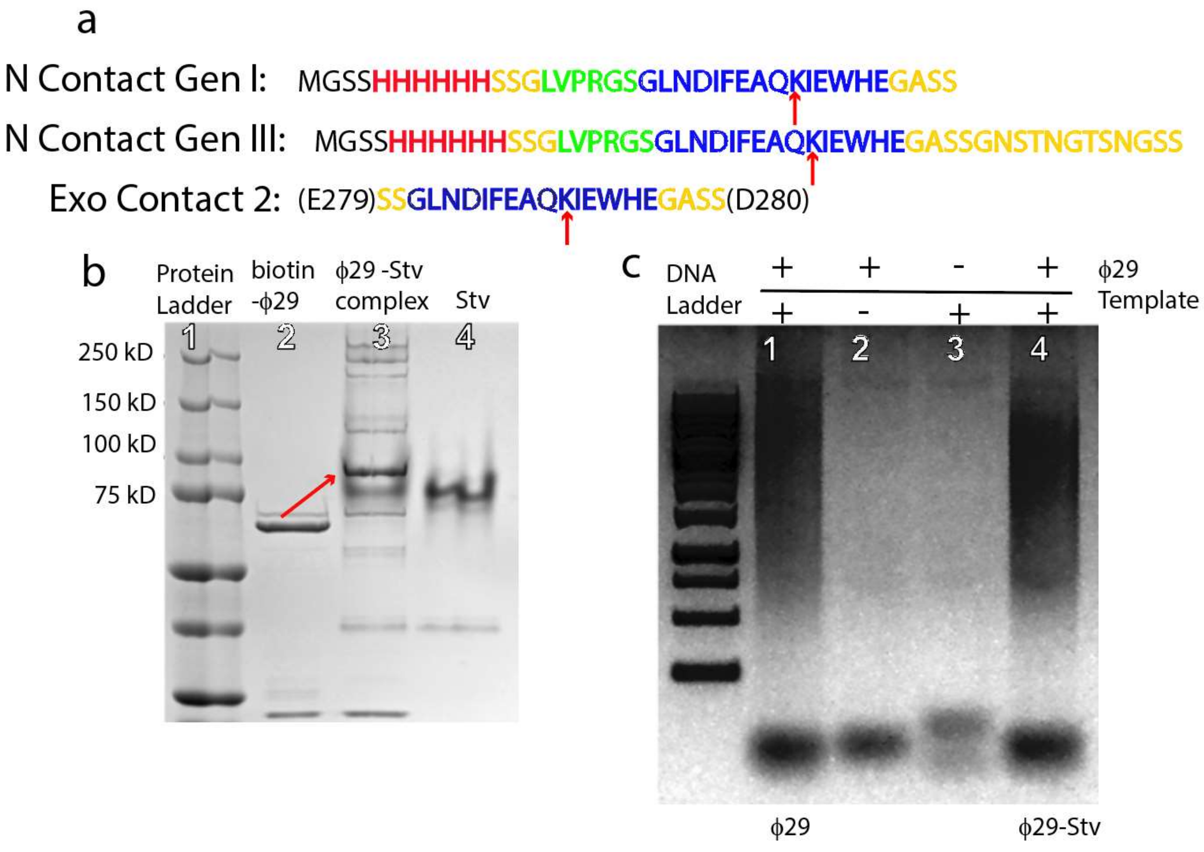


**Figure S1 Protein sequences, synthesis and gel analysis**.

(a) Avitag sequence (blue) inserted at the N terminus (N Contact Gen 1). The epsilon-amine of the central lysine K (marked by the red arrow) is the target of biotinylation by the BirA enzyme. Linking sequences are marked in yellow, green is a thrombin cut site and red marks a His tag. N Contact Gen III lists the sequence inserted at the N terminus in the third generation polymerase incorporating an additional length of flexible linker. Generations II and III incorporate a second avitag (Exo Contact 2) in the exonuclease domain between E279 and D280. The polymerase is rendered exonuclease-deficient with D12A and D66A mutations and was synthesized as follows: *E. coli* strain BL21 (DE3) (Novagen) was transformed with different versions of the φ29 gene embedded in pET15b plasmid and grown on an LB agar plate (1% tryptone, 0.5% yeast extract, 0.5% NaCl, and 1.5% agar) containing ampicillin (50 µg/ml) to select the transformants. Cells were grown in LB medium (20 ml) at 37°C with shaking for 12 hours. A part of the culture was diluted 1:1000 into fresh LB medium (1L) and grown at 37°C with shaking. The 1L culture was induced by adding 0.5 mM IPTG when the $OD_{600}$ reached 0.6 and kept shaking overnight at 18°C. Cells were harvested in a 1L centrifuge bottle by centrifugation at 5,710g for 20 minutes at 4°C and stored at -80°C until required. The purification is essentially as described elsewhere.[1] For subsequent BirA biotinylation, the purified protein was exchanged to a buffer containing 20 mM potassium phosphate buffer pH 7.0, 200 mM L-Glutamic acid potassium salt and 1 mM DTT. The *in vitro* enzymatic biotinylation was performed by incubating 100 µg of polymerase in the same buffer with 10 mM ATP, 10 mM $Mg(OAc)_2$, 50 mM biotin and 15 units of BirA (Avidity) for 1 hour at 30°C. Free biotin was removed by a desalting column (GE Life Sciences).

(b) The formation of a complex with streptavidin was verified by a protein gel which shows how the biotinylated Φ29 (lane 2) forms a complex with streptavidin molecule (lane 3). Lane 4 is streptavidin alone. Higher molecular weight bands in lane 3 are complexes incorporating more than one polymerase.

(c) Activity of modified polymerase complexed with streptavidin: A linear single stranded oligonucleotides RCR (5'- p -CCGTACGATTCGTATCTACTATCGTTCGATTCGCATCATCTA -3') was used to form circular RCR template by enzymatic self-ligation with Circligase (Epicentre). 0.1 nmol linear single strand RCR DNA was mixed with 100 Units Circligase in 1 X reaction buffer containing 50 µM ATP and 2.5 mM $MnCl_2$. After 2 hours incubation at 60°C, the product was heated up to 80°C for 10 minutes to inactivate the Circligase. The linear ssDNA left in the solution was digested by Exo I (NEB). The RCR template was analyzed by electrophoresis on a denaturing gel containing 8 M urea and 20% polyacrylamide for quality control. 2.5 pmols of RCR template was annealed with 50 pmols RCR primer (5'-GGCATGCTAAGCATAGATGAT -3') by heating up to 95°C for 5 minutes and gradually cooling down to room temperature (decreasing 0.1°C/s) and stored at -20°C for later use. A rolling circle replication reaction was performed for the activity test of all versions of φ29 DNA polymerase. 1.25 pmols RCR template and primer complex was mixed with 500 µM dNTP and 4 pmols φ29 DNA polymerase in 1 X reaction buffer containing 50 mM Tris-HCl ph7.5, 10 mM $MgCl_2$, 10 mM $(NH_4)_2SO_4$, 4 mM DTT. The mixture was incubated at 30°C for 1 hour. The product was visualized on 0.8% agarose gel by GelRed (Biotium) staining. Polymerization products of the modified Φ29 are shown in lane 1. The production of polymers is essentially unaltered when the Φ29 is complexed with streptavidin (lane 4). The assays were repeated in the buffer used for STM measurments (1mM phosphate buffer, pH=7.4, 4 mM TCEP, 10mM $MgCl_2$ with 1 mM dNTPs and 1 µM template) to ensure that activity was maintained.

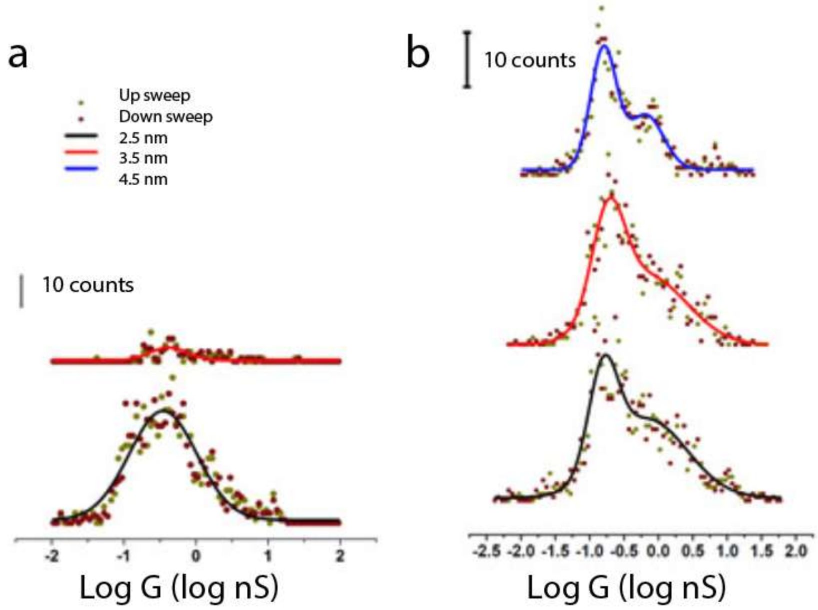


**Figure S2:** Conductance distributions as a function of gap size for (a) streptavidin-functionalized electrodes and (b) after the introduction of Gen I monobiotinylated Φ29.

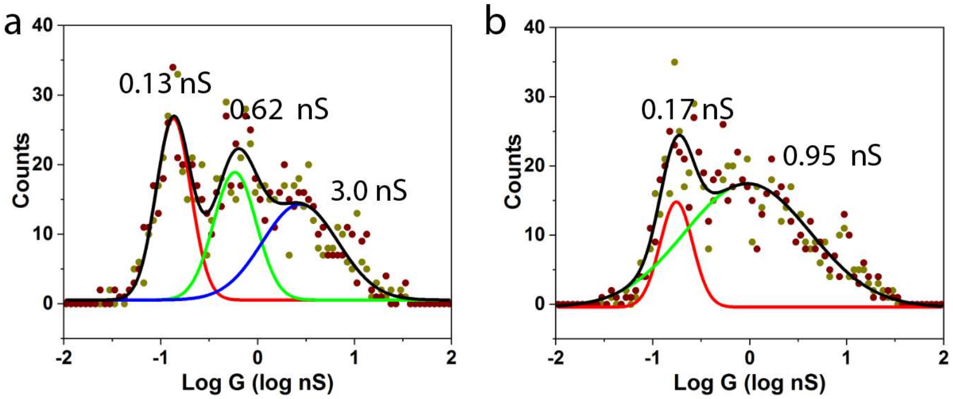


**Figure S3:** (a) Conductance distribution for the Gen II bisbiotinylated polymerase contacted via thiolated streptavidin to the electrodes (Gap = 4.5 nm). Note that the highest conductance peak is smaller than that observed for wild-type streptavidin connected to the electrodes via thio-biotin (Figure 1f), consistent with observations of the conductance of streptavidin alone. [2] (b) Conductance distribution for the Gen II polymerase attached to a thiolated substrate and contacted with a bare probe, indicating that the two smallest peaks in the conductance distribution arise from interactions between surface thiols on the Φ29 and the bare metal.

a

5'CCCCCCCCCC AACTGGCCG TCGTTTTACA TATGTAAAC GACGGCCAGT T3'

```
┌ACATTTTGCTGCCGGTCAACCCCCCCCCC 5′
T    |||||||||||||||
└ATGTAAACGACGGCCAGTT
```

b

5'AAAAAAAAAA AACTGGCCG TCGTTTTACA TATGTAAAC GACGGCCAGT T3'

```
┌ACATTTTGCTGCCGGTCAAAAAAAAAAAA 5′
T    |||||||||||||||
└ATGTAAACGACGGCCAGTT
```

c

5'CCCCCCCCCC AAAAAAAAAA AACTGGCCG TCGTTTTACA TATGTAAAC GACGGCCAGT T3'

```
┌ACATTTTGCTGCCGGTCAAAAAAAAAAAACCCCCCCCCC 5′
T    |||||||||||||||
└ATGTAAACGACGGCCAGTT
```

d

5'ATC ATC ATC ATC ATC AACTGGCCG TCGTTTTACA TATGTAAAC GACGGCCAGT T3'

```
┌ACATTTTGCTGCCGGTCAACTACTACTACTACTA 5′
T    |||||||||||||||
└ATGTAAACGACGGCCAGTT
```

**Figure S4:** Sequences of the DNA templates used and their folded structures

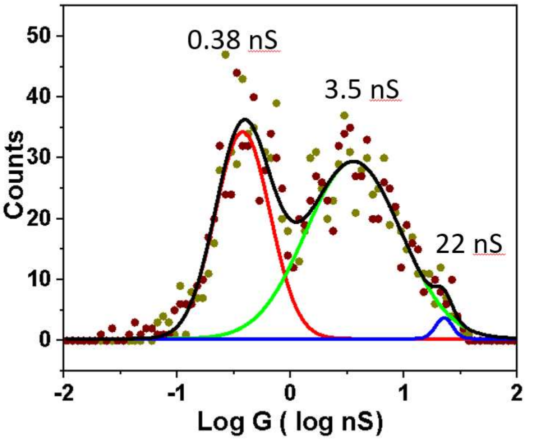


**Figure S5:** Conductance distribution for Φ29 in the closed form - repeat measurement.

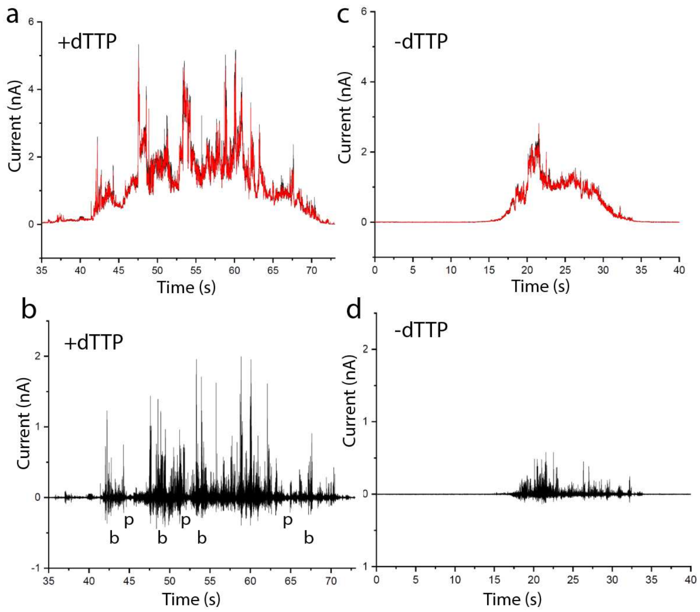


**Figure S6: Characterizing the noise signals generated when polymerase is activated.** (a) The current through the polymerase changes markedly over time as the STM probe drifts (mainly obscured black curve). This drifting baseline current is fitted with an asymmetric least squares (ALS) procedure to yield a smoothed background current (red curve superimposed on black curve). (b) Subtraction of the fitted background shows the rapid changes in current that occur in an activated polymerase. Typically the dynamic signals occur in bursts (b) interspersed with pauses (p). (c) Shows the current recorded vs time in the absence of the complementary nucleotide triphosphate, and the ALS fit. These data were chosen to have about the same DC conductance as the active polymerase in (a) and (b). (d) The baseline subtracted signal shows that there is also noise in the inactive system.

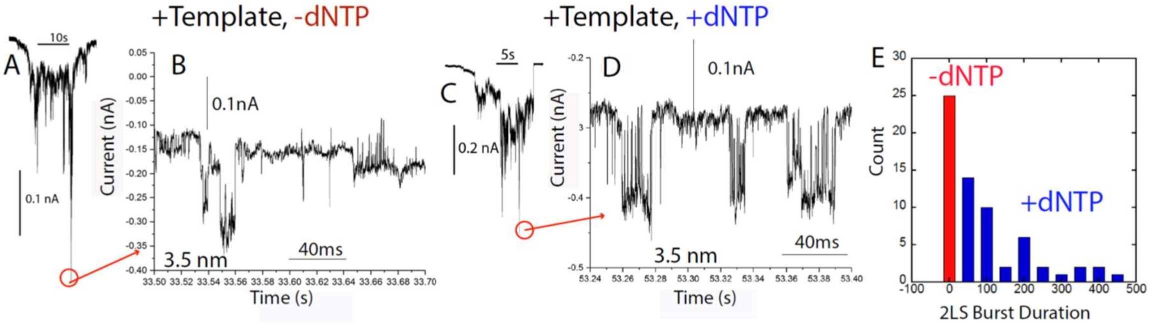


**Figure S7:** Noise signals obtained with the Gen I monobiotinylated polymerase. (A) Current vs time with bound template but no dNTPs. The highest current region is expanded in (B). There are many jumps in the baseline but no obvious two-level telegraph noise. (C) Current-time recording with dNTPs added. Noise spikes are more evident and, when an expanded trace is plotted (D) obvious bursts of two-level random telegraph noise are present. (E) is a histogram of burst duration, clearly longer for the +dNTP case. Note: in these plots current is increasing in the downward direction.

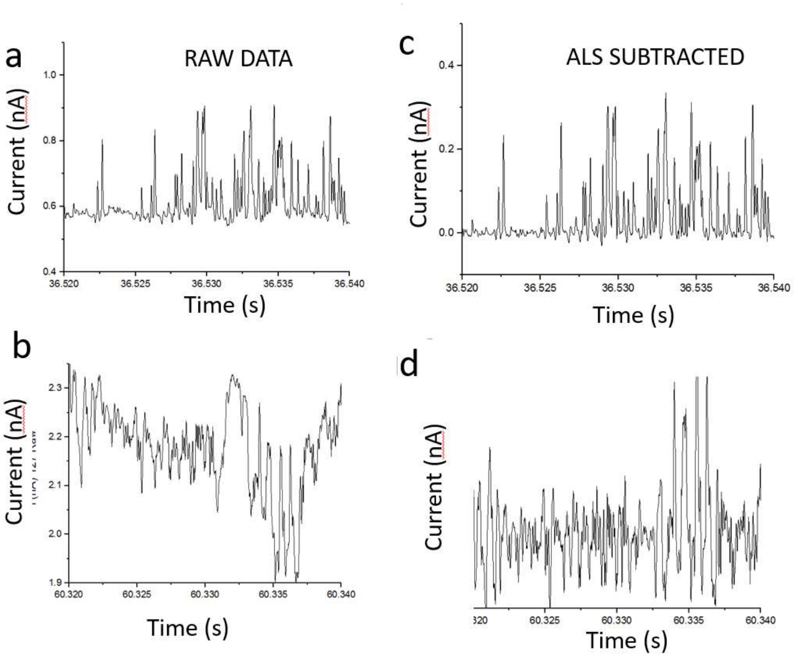


**Figure S8:** Effect of ALS background subtraction (smoothing factor = 0.1 ms). (a) and (c) are raw and subtracted data for a region of relatively flat baseline. (b) (raw) and (d) (subtracted) illustrate how the noise features are well-preserved even in the presence of large baseline variations.

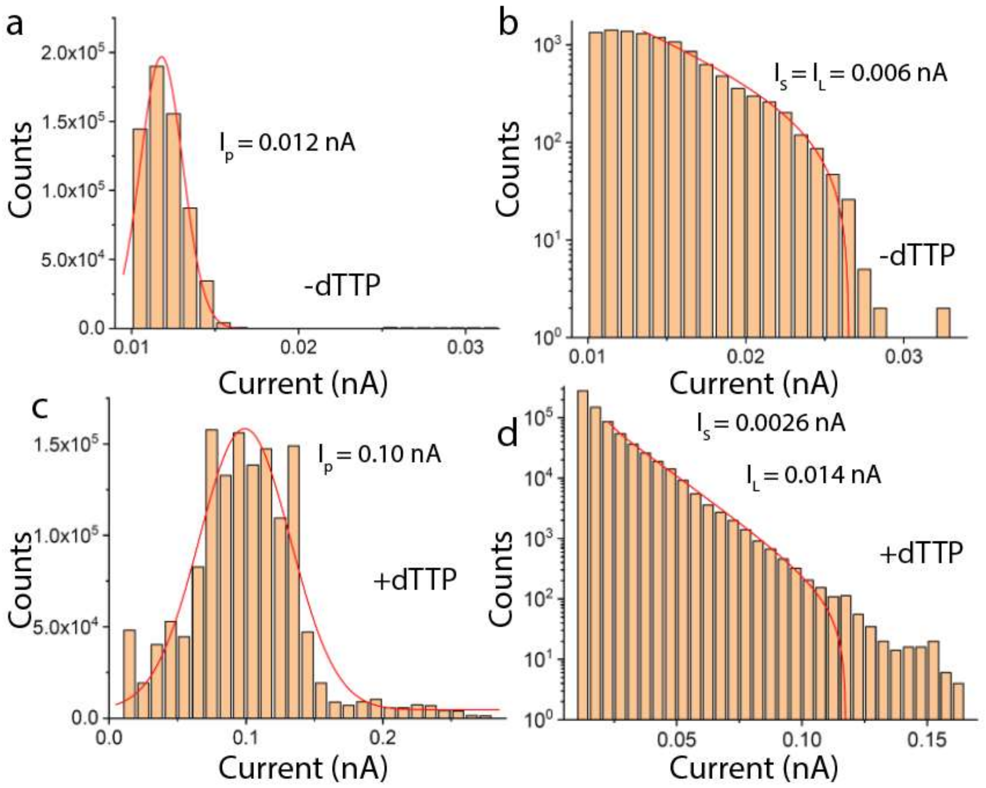


**Figure S9: Small and large fluctuations defined in terms of baseline current.** Larger signals are obtained with the higher-conductance contacts. To quantify this, the baseline levels in a given run are characterized by fitting a Gaussian model to the distribution of measured currents, characterized by the peak value of the Gaussian distribution, $I_P$ (a, c). A two exponential fit ($I_{S,}$ $I_L$) is used to model the distribution of noise signal amplitudes. When the complementary base is absent (b) the fit converges to one value, $I_S = I_L$ in all cases. (d) In the presence of the complementary nucleotide, 9 out of 13 molecules showed a bimodal distribution of spike heights with $I_S << I_L$.

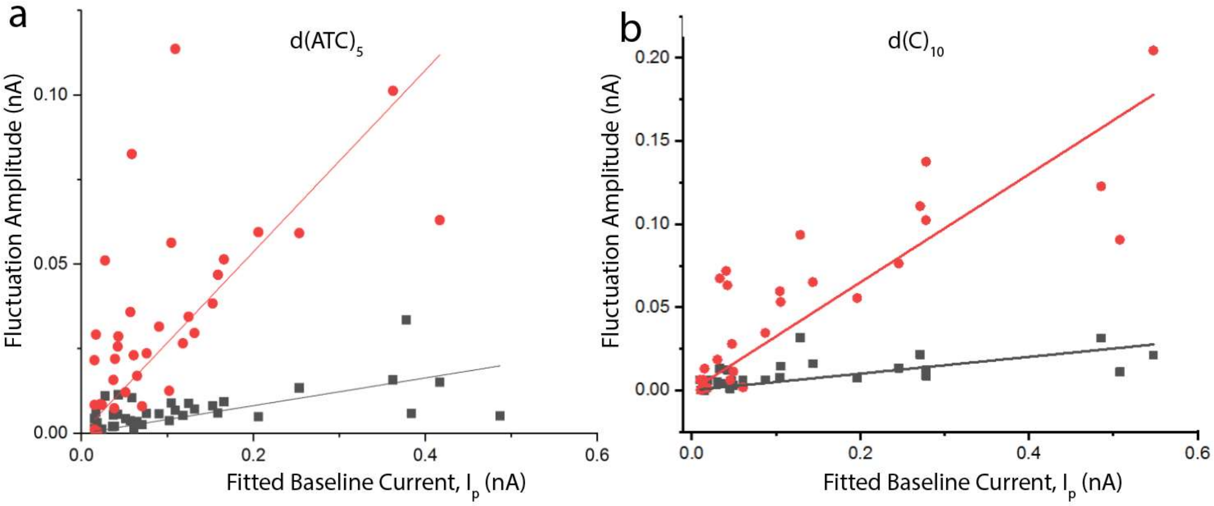


**Figure S10:** Values of $I_L$ (red) and $I_S$ (gray) plotted vs the associated value of baseline current ($I_P$). for 38 Φ29 molecules actively transcribing the $d(ATC)_5$ template (a) and 25 molecules transcribing the $dC_{10}$ template (b).

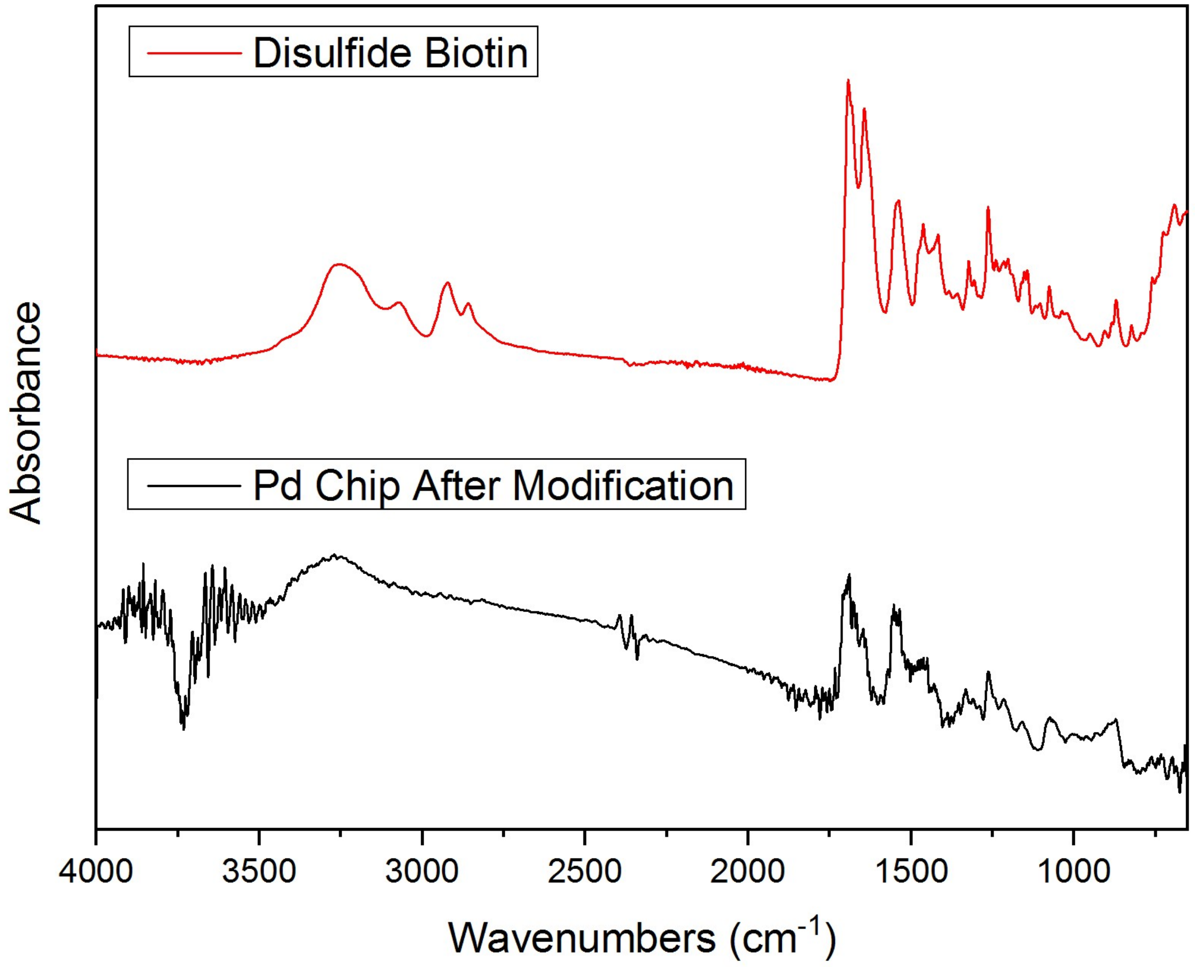


**Figure S11 FTIR** scans showing biotin functionalization of Pd.

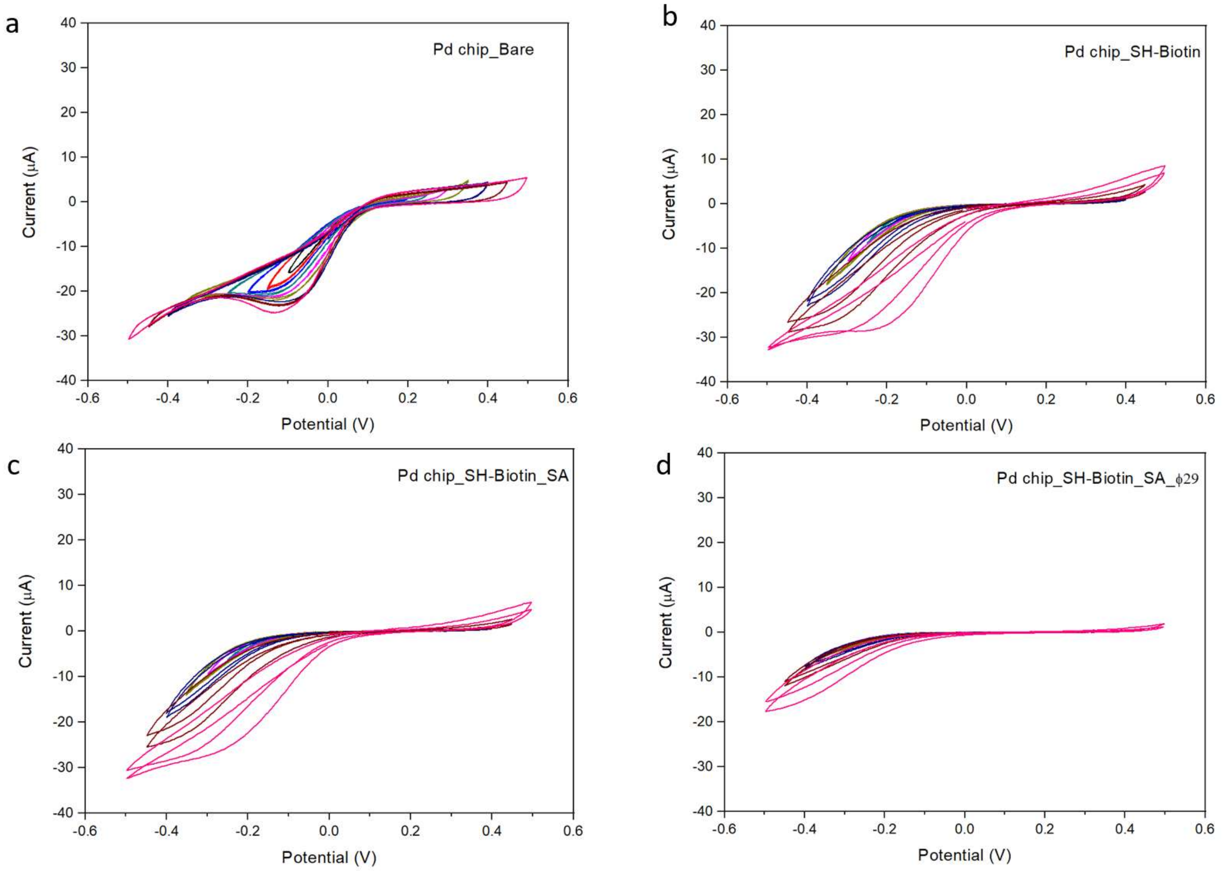


**Figure S12 CV scans of functionalized Pd.** Pd substrates were cut into 0.5 cm x 4.0 cm in size and used as the working electrode, with an active cell area of about 0.5 cm x 1.0 cm. The substrate was treated with a hydrogen flame before functionalization. Cyclic voltammetry was performed on a potentiostat (Model AFCBP1, Pine Instruments), using a Pt wire as the counter electrode and an Ag/AgCl (3M KCl) as the reference electrode. The maximum sweep range is from -0.5 V to +0.5 V, with a sweep rate of 10 mV/s, starting from ±0.05 V and increasing the scan range on 0.05V increments. (a) Bare Pd. (b) Functionalized with SH-Biotin (50 μM ,overnight). (c) With streptavidin added (1μM, 0.5 h). (d) And after addition of Gen III Φ29 polymerase (1μM, 2h). Two repeated sweeps are shown at the highest bias, showing some instability in the adsorbate.

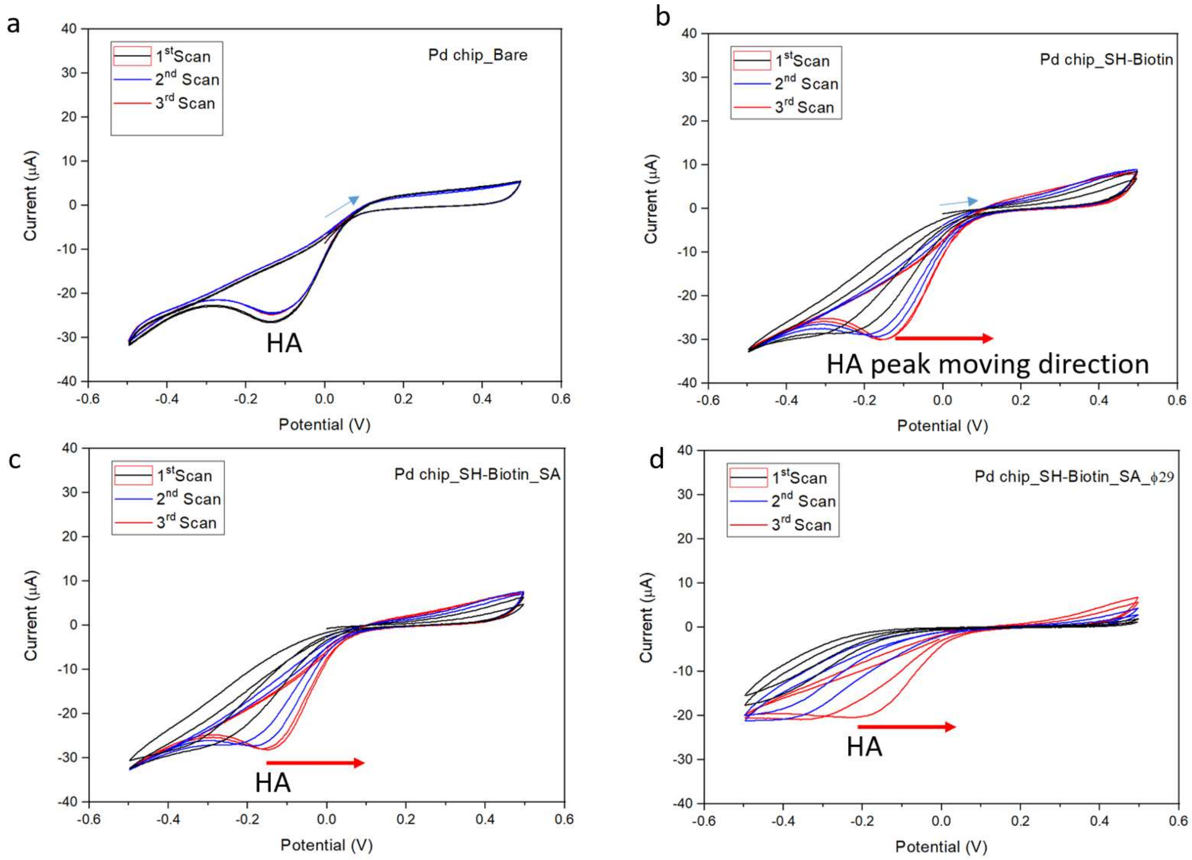


**Figure S13 Sweeps to negative potentials destabilize thiol-bound adsorbates.** (a) Repeated sweeps on bare Pd reproduce well (HA is the hydrogen adsorption peak). On functionalization with biotin (b), biotin plus streptavidin (c) and biotin plus streptavidin plus Φ29 polymerase, the HA peak, initially moved to more negative potentials relative to bare Pd, moves up towards the potential observed in bare Pd, consistent with stripping of the adsorbate by thiol reduction.

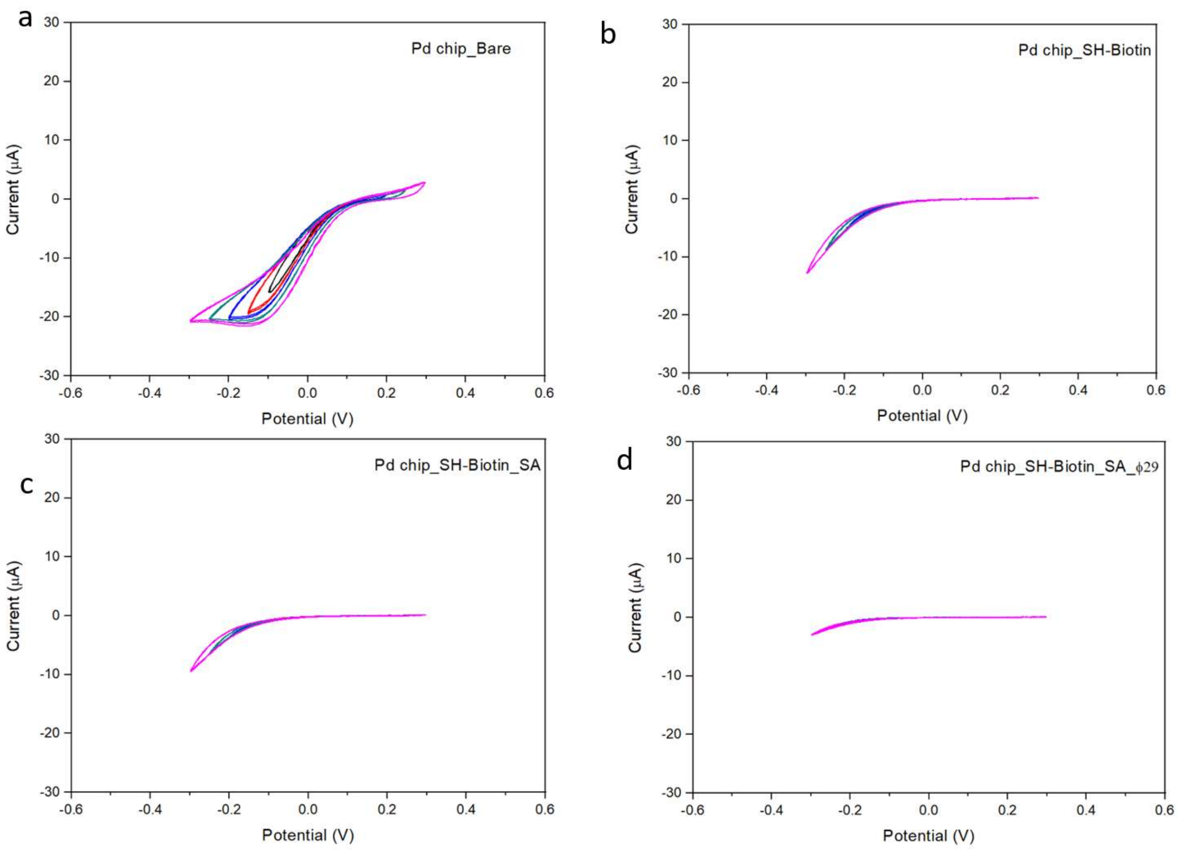


**Figure S14 Electrodes are passivated by biomolecular adsorbates in the potential range used for electronic measurements.** (a) Repeated sweeps on bare Pd. On functionalization with biotin (b), biotin plus streptavidin (c) and biotin plus streptavidin plus Φ29 polymerase the electrode becomes increasingly passivated. These measurements illustrate that these electronically conductive films do not exchange charge with ions in solution.
